# Towards Consensus Gene Ages

**DOI:** 10.1101/042036

**Authors:** Benjamin J. Liebeskind, Claire D. McWhite, Edward M. Marcotte

## Abstract

Correctly estimating the age of a gene or gene family is important for a variety of fields, including molecular evolution, comparative genomics, and phylogenetics, and increasingly for systems biology and disease genetics. However, most studies use only a point estimate of a gene’s age, neglecting the substantial uncertainty involved in this estimation. Here, we characterize this uncertainty by investigating the effect of algorithm choice on gene-age inference and calculate consensus gene ages with attendant error distributions for a variety of model eukaryotes. We use thirteen orthology inference algorithms to create gene-age datasets and then characterize the error around each age-call on a per-gene and per-algorithm basis. Systematic error was found to be a large factor in estimating gene age, suggesting that simple consensus algorithms are not enough to give a reliable point estimate. We also found that different sources of error can affect downstream analyses, such as gene ontology enrichment. Our consensus gene-age datasets, with associated error terms, are made fully available at so that researchers can propagate this uncertainty through their analyses (https://github.com/marcottelab/Gene-Ages).

## Introduction

From their inception, whole genome datasets have been used to infer the evolutionary history of gene families [1]. The age of a gene family, its provenance, and its evolutionary history, such as loss and duplication events, can inform us about its function [2]. For instance, gene age has been found to correlate with disease-association [3,4], evolutionary rate [5], and the number of associated protein-interaction partners [6], and a gene’s phylogenetic distribution can be used to infer aspects of its function [7]. Gene-ages can also be used to estimate the gene content of ancient organisms, such as the last universal common ancestor (LUCA, [1]), or the last eukaryotic common ancestor (LECA, [8,9]). Accordingly, an analysis of gene family ages on a genomic scale can inform the phylogenetic history of important phenotypes, such as eyes or the nervous system [10,11]. In more recent years, gene age has been used to annotate systems biology datasets [12–14], with the promise of elucidating the evolutionary history of core cellular machinery.

Such studies rely first and foremost upon the correct identification of homologs and/or orthologs. These two relationships form the basis of the gene-age determination in nearly all studies, with orthology being the more common criterion [3,15]. Orthology is a pairwise relationship between two genes that occurs when their most recent common ancestor (MRCA) lies at a speciation event in a phylogenetic gene tree. This is in contrast to paralogs, whose MRCA lies at a gene duplication event (nodes on gene trees represent either speciation or gene duplication events, barring horizontal gene transfer)[16,17]. Orthologs tend to display higher functional conservation than paralogs (though perhaps only weakly [18] - see [19] for a review), hence their use as a basis of cross-species comparison. Typically, studies of gene age will consider an orthologous group to be all the descendent lineages of the deepest speciation node, or the divergence between the two most distant homologs, if that is the criterion being used, as in “phylostratigraphy” [4]. Then, the age of the gene group is defined as the MRCA of the species found in that group.

Inferring a gene family’s age thus relies on the accuracy of orthology assignment, but inferring correct orthologs is notoriously difficult, with no one of the more than 30 algorithms out-performing all others [20]. In particular, algorithms differ strongly in the tradeoff between recall and precision [20]. Yet most studies on gene age rely on only one kind of algorithm, either using a pre-existing method or establishing an *ad hoc* protocol, most of which resemble one of the pre-existing algorithms [3]. Although methods for probabilistic orthology assignment do exist [21], available methods are not currently scalable to large genomic datasets using protein sequences, and at any rate still rely on a preliminary clustering step to infer gene families. Consensus algorithms also exist, some of which seem to substantially improve performance on established benchmarks [22,23]. However, these methods still give only a point estimate. Another approach is to propagate the uncertainty that necessarily arises in orthology inference through subsequent analyses. However, it is unclear what the relevant sources of uncertainty are in orthology inference, and most consensus algorithms do not keep track of the different sources of error.

To remedy this situation, we characterized the error structure of gene-age estimation using 13 popular orthology inference algorithms. In doing so, we identify common types of errors and, after correcting these, present consensus gene-age calls for several model organisms (Table 1). We provide these gene-age estimates along with a detailed analysis and annotation of the uncertainty associated with each age call so that this uncertainty can be propagated through future analyses, as we show for functional term enrichment. The consensus gene ages we calculate can be used for annotating genomic datasets in a variety of fields, and the analysis of error will help prioritize genes for manual annotation and aspects of orthology inference for future study.

**Table 1.** Species for which final consensus tables were constructed. Tables are available at geneages.org.

| Common Name | Uniprot ID | False-negative analysis |
| --- | --- | --- |
| Anopheles gambiae (Mosquito) | ANOGA | No |
| Bos taurus (Cattle) | BOVIN | No |
| Branchiostoma floridae (Lancelet) | BRAFL | No |
| Caenorhabditis elegans (Worm) | CAEEL | Yes |
| Candida albicans | CANAL | Yes |
| Canis lupus familiaris (Dog) | CANFA | No |
| Gallus gallus (Chicken) | CHICK | Yes |
| Ciona intestinalis (Tunicate) | CIOIN | No |
| Cryptococcus neoformans | CRYNJ | No |
| Danio rerio (Zebrafish) | DANRE | Yes |
| Drosophila melanogaster (Fly) | DROME | Yes |
| Homo sapiens (Human) | HUMAN | Yes |
| Ixodes scapularis (Tick) | IXOSC | No |
| Macaca mulatta (Rhesus macaque) | MACMU | No |
| Monosiga brevicollis (Choanoflagellate) | MONBE | No |
| Monodelphis domestica (Opossum) | MONDO | No |
| Mus musculus (Mouse) | MOUSE | Yes |
| Nematostella vectensis (Sea anemone) | NEMVE | No |
| Neurospora crassa | NEUCR | No |
| Ornithorhynchus anatinus (Platypus) | ORNAN | No |
| Pan troglodytes (Chimp) | PANTR | No |
| Phaeosphaeria nodorum | PHANO | No |
| Rattus rattus (Rat) | RAT | Yes |
| Saccaromyces cerevisiae (Budding yeast) | YEAST | Yes |
| Schistosoma mansoni (Blood fluke) | SCHMA | No |
| Schizosaccharomyces pombe (Fission yeast) | SCHPO | Yes |
| Sclerotinia sclerotiorum | SCLS1 | No |
| Takifugu rubripes (Pufferfish) | TAKRU | No |
| Ustilago maydis (Corn smut/Huitlacoche) | USTMA | No |
| Xenopus tropicalis (Frog) | XENTR | No |
| Yarrowia lipolytica | YARLI | No |

## Results

### Data collection

In order to fairly consider the range of orthology algorithms, we took advantage of the reference datasets managed by the Quest for Orthologs (QFO) consortium. QFO researchers have established community standards and benchmarks for orthology inference and have made their benchmarking results publicly available [20]. Importantly, the algorithms that have contributed to the benchmarking tool are widely used and capture the variety of methods commonly used in the literature to infer orthology and gene age [24–32]. It is therefore expected that nearly every study of gene age, regardless of the method used, will closely resemble the results of at least one of the algorithms we explore here. We downloaded orthology calls for 66 reference proteomes based on 13 orthology inference algorithms from the QFO website and inferred the ages for each human gene by mapping the species in each ortholog group onto a reference species tree from SwissTree, which was derived from a consensus of trees found in the literature [33]. The results below are with reference to the human proteome, but the same methods were applied to a variety of model organism proteomes (Table 1).

### The effect of algorithm choice on the distribution of human gene ages

To investigate the effect of algorithm choice on inferred gene age, we broke the reference species tree into eight age categories (Figure 1). These categories form nested clades, with the exception of the category “Euk+Bacteria.” This non-phylogenetic category captures the substantial number of eukaryotic genes that were horizontally transferred from bacteria after eukaryotes diverged from the rest of archaea [34,35], and is defined as genes present in eukaryotes and bacteria but not archaea. For each algorithm, every human gene was assigned to the age category in which the MRCA of the species in its orthogroup falls, and the distributions over the different age categories for the human proteome inferred by that algorithm was calculated.

**Figure 1.**
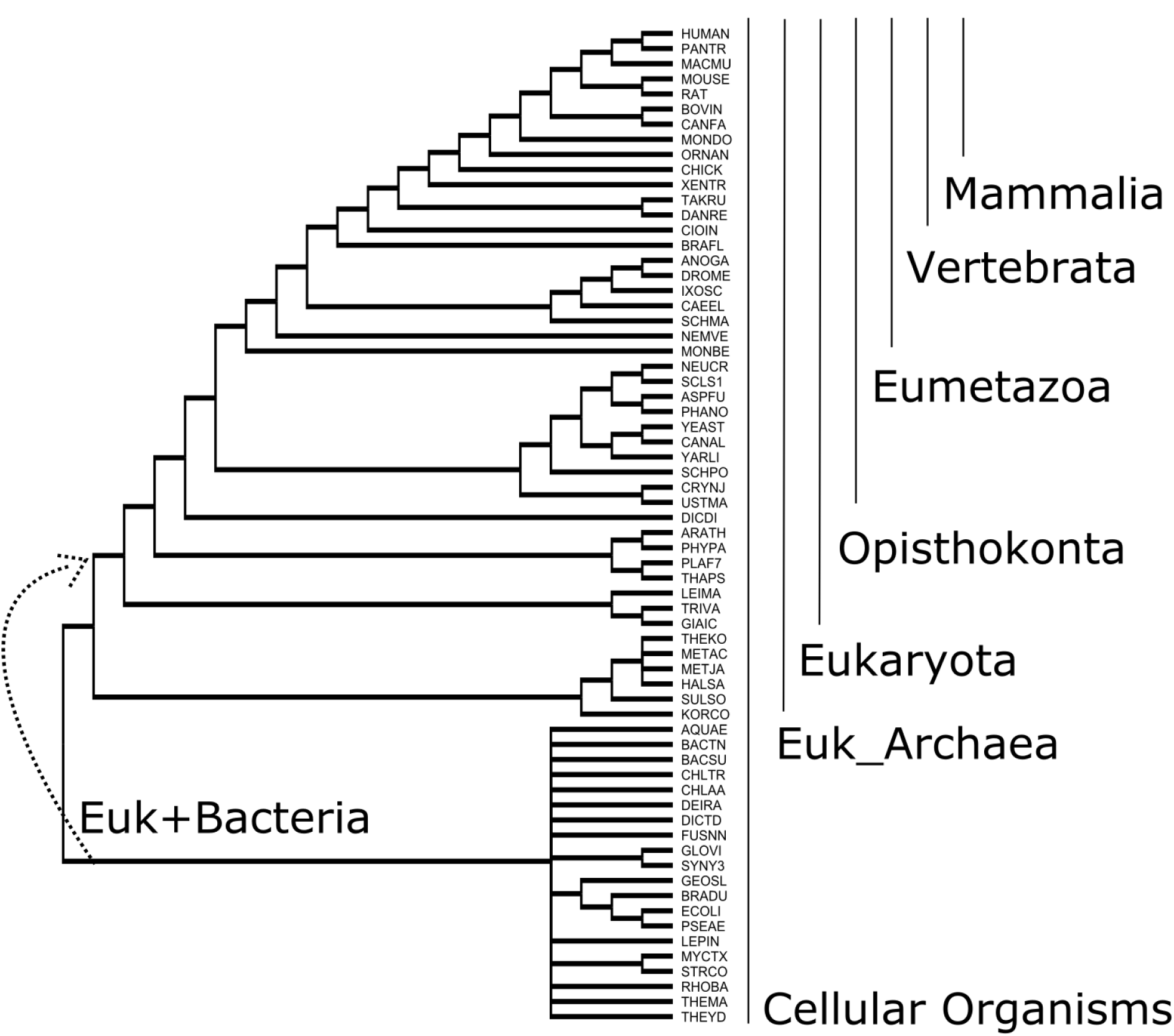
The reference species tree and age categories used for gene-age inference. This tree is based on SwissTree [33] and reflects a consensus of recent large-scale phylogenies. Tip names are Uniprot species identifiers.

We found that the algorithms fell into two distinct groups with respect to the distribution of age classes. Clustering the algorithms by the average patristic distance between their per-gene age calls recapitulated this grouping (Figure 2), and we define the two groups based on the midpoint root of this tree. One group tended to find that most orthogroups could be traced to the MRCA of vertebrates, whereas the other group found a much older mode age dating back to LECA. We call these two groups the “young” and the “old” group respectively, though of course there are many more subtle and interesting distinctions between the algorithms.

**Figure 2.**
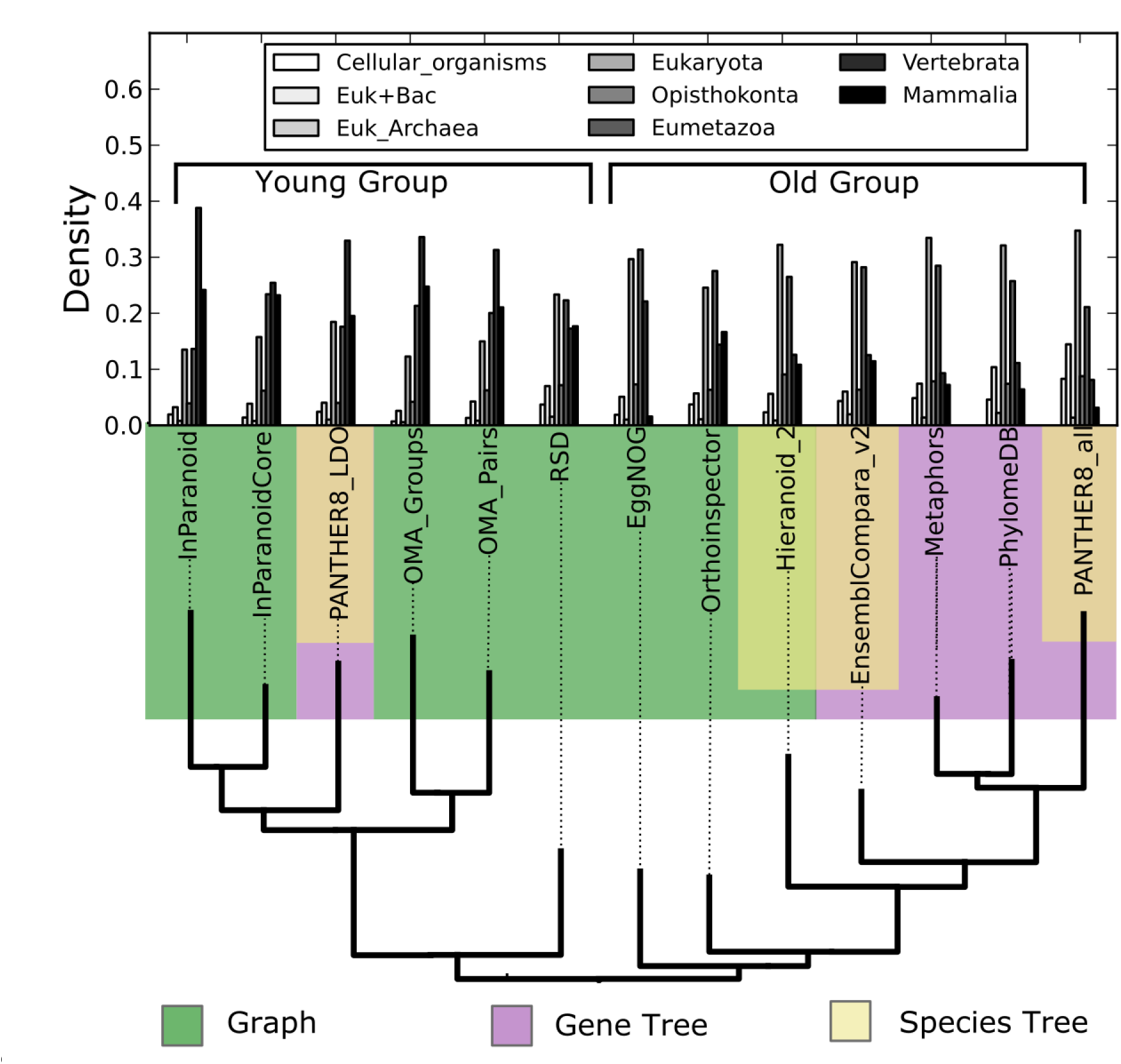
Distribution of age categories in the human proteome inferred by 13 different orthology inference algorithms. Algorithms were clustered according to the average pairwise distance between their age-calls, counted in units of braches (patristic distance). The distance tree is rooted at the midpoint. Algorithms are colored by the methods they use to infer orthology. They either use a graph-based or a gene tree-based strategy, either with, or without, the use of a species tree.

Orthology inference algorithms are typically classed into graph-based and tree-based methods [20]. However, we found that even though tree-based methods tended to fall in the “old” group, this was not universally the case, nor were all graph-based methods found in the “young” group. The use of species tree information was not a determining factor either (Figure 2). The bimodal nature of the age calls, either “young” or “old”, is therefore not simply a reflection of the graph/tree distinction, although it is clearly correlated. What is the source of this bimodality? One obvious answer is systematic error in the “young” group algorithms, the “old” group, or both. Systematic error in the young group would be equivalent to false negatives, i.e. missing orthology assignments, whereas systematic error in the old group is equivalent to false positives, or spurious orthology assignments. This would have the effect of pushing the age of the group away from or towards the root of the tree, respectively.

### Identifying systematic error

We first investigated whether the bimodality of age-calls played out on the single gene level or whether the two groups apparent in Figure 2 were due to the effects of averaging across genes, with error being randomly distributed among proteins. To do so, we calculated a simple statistic that captured how bimodal a protein’s age calls were between the two groups of algorithms (“old” and “young”). This statistic, which we call bimodality, is the difference between age-call variation within the two groups and between them, with more highly bimodal proteins having more variation between groups. Over 80% of proteins had some degree of bimodality corresponding to these two age groups, or none, as is expected given the hierarchical clustering in Figure 1. The remaining genes were anti-correlated with the “old”/”young” groupings. Furthermore, the degree of bimodality between the “young” and “old” algorithm groups correlates well with the amount of error associated with each protein (Spearman’s *ρ*: .65)(Figure 3). That is, proteins with a large amount of error tend to be more bimodal. The bimodality between algorithms is therefore a systematic phenomenon and a major source of error in these datasets. Unfortunately, in the case of highly polarized genes, we cannot know *a priori* whether the “old” or “young” age is the correct one. It is therefore important to propagate this uncertainty through further analyses, and the bimodality statistic is included with our consensus age estimates.

**Figure 3.**
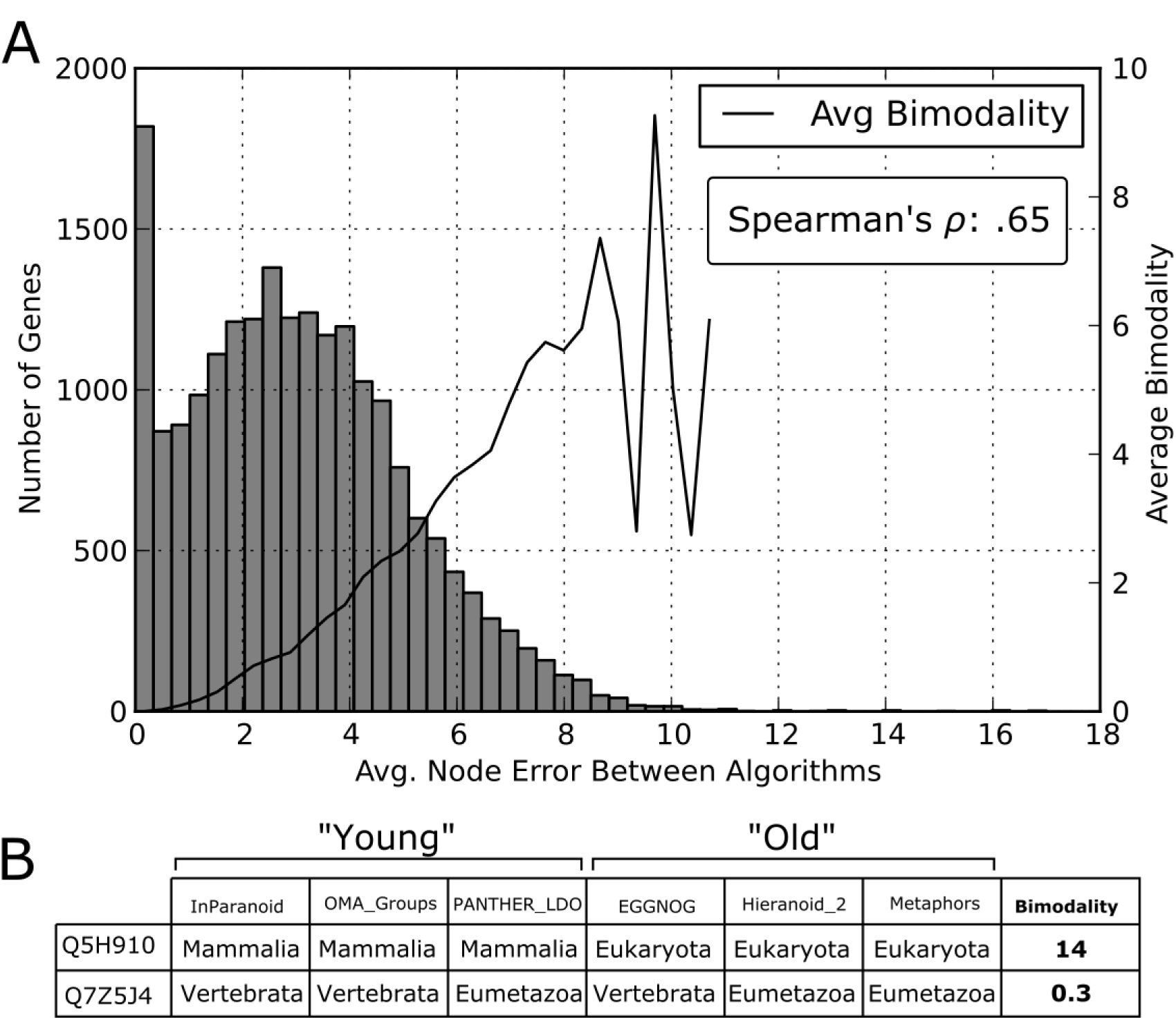
Error statistics. (A) The distribution of average node error, a measure of disagreement among the algorithms for a given gene, is given, along with a plot of the average bimodaliy in each bin. Genes with more error tend to be more bimodal between “old” and “young” algorithms. (B) Example of a strongly bimodal and weakly bimodal gene with a few representative algorithms. The ages are given as categories for clarity, but the bimodality statistic is calculated according to patristic distance between node age-calls (Methods).

We also investigated whether aspects of the individual proteins contributed to systematic error. For instance, it may be difficult to infer correct evolutionary relationships for small proteins, or those with many domains. At least one orthology inference algorithm uses this idea to “correct” for protein length [36]. However, we found that protein length has a weak positive correlation with age-call error, and that the number of domains also correlates weakly (Spearman’s *ρ*: < .2 in both cases).

### Systematic false negatives

What are the causes of systematic false negatives and can we identify them without *a priori* knowledge of the true orthogroup? One clue comes from the different age-category distributions between PANTHER8_all and PANTHER8_LDO [25]. These two sets of orthology calls are based on the same set of gene trees, but differ in their definition of orthology. “LDO” stands for “least diverged ortholog,” and only considers the least diverged among a set of co-orthologs to be the true ortholog of an outgroup. This can be contrasted to the traditional phylogenetic definition of orthology where all co-orthologs are equally orthologous to the outgroup (Figure 4) [17]. Although it may be useful to split co-orthologous groups, as the LDO definition does, in cases where orthology is being used for, e.g. gene function annotation, it is inappropriate for defining the age of a gene or gene family because the age must be in reference to the topology of the phylogenetic tree. The fact that PANTHER8_LDO’s age category distribution resembled that of several graph-based methods, and the fact that it clustered with them based on its per-gene age calls (Figure 2), suggests that these methods may be splitting up co-orthologous groups as well.

**Figure 4.**
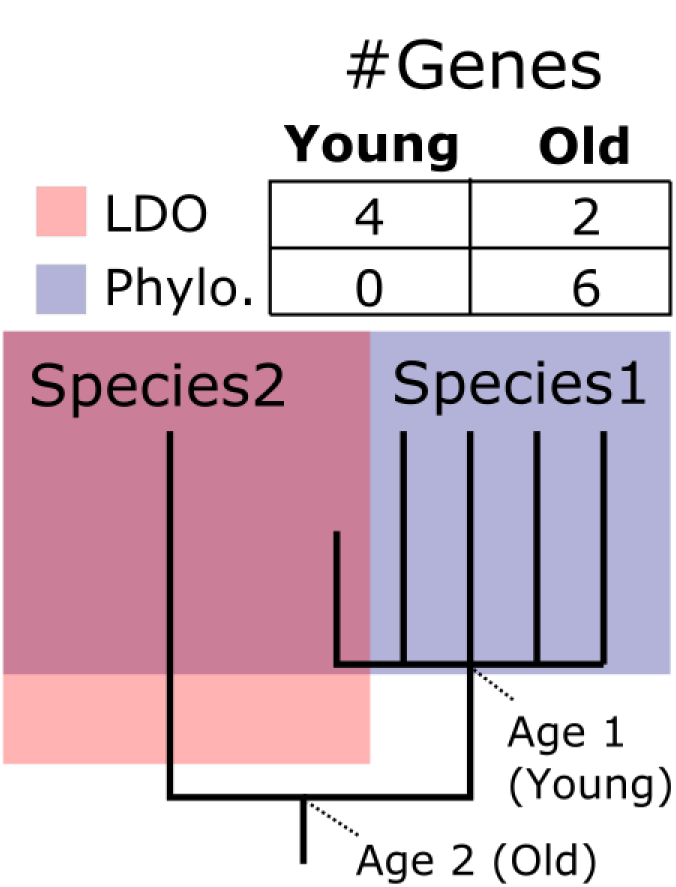
Determination of false negatives due to co-ortholog over-splitting. This tree compares the ages given by least derived orthology (LDO) and traditional, phylogenetic orthology (Phylo.). Given a group of co-orthologs in Species 1, LDO will give only the co-ortholog with the shortest distance to an outgroup (gene in Species 2) the status of ortholog to this outgroup (red box). All others are put in separate orthogroups. Hence, LDO produces more genes that are mapped (incorrectly) to a younger age (**Y**), whereas traditional, phylogenetic orthology (blue box) includes all co-orthologs to the orthogroup, thereby mapping more genes to the older age (**O**).

There is no gold standard set of co-orthologs in this dataset, so we used the database PhylomeDB as a reference for identifying co-ortholog over-splitting. To do so, we downloaded PhylomeDB summary files for 10 species in PhylomeDB’s model species collection (PhyC2) that overlapped with species in our tree (Table 1), and determined groups of co-orthologs that were then used for the analysis. Briefly, for protein (A), if an algorithm called a younger age (Y) and PhylomeDB an older age (O), and if in the co-orthologs of (A) we could find a protein (B) which that algorithm called at age (O), then (B) was identified as the LDO, age (O) was assumed to be the true age, and that algorithm was determined to be over-splitting the co-ortholog group (Figure 4). This was not carried out for proteins on which PhylomeDB’s age call was determined to be a false positive (see below). We note that this method for identifying co-ortholog over-splitting is not ideal, because it relies on a single, imperfect algorithm (PhylomeDB). It is conservative, however, because algorithms will only be trimmed if they give a member of the co-orthologs the *exact* same age on the species tree as that called by PhylomeDB on the focal gene. More thorough analyses of whether graph-based methods are consistently missing co-orthologs will be necessary in the future.

### Identifying false positives

If genes of distant organisms are incorrectly inferred to be part of an orthology group, it will drive the age of the orthogroup towards the root of the tree. Recent HGT events are a biological source of such errors, but some algorithmic error is expected to play a role as well. Such problems are perhaps more likely to occur in tree-based algorithms, where slight re-arrangements that don’t strongly affect the likelihood of the tree can have an outsized effect on the inference of gene gains and losses [37]. In such cases, the large number of taxa that fall between the true in-group taxa and the false positive out-group taxa will be inferred to have lost the orthogroup. We used this criterion on a per-gene basis to identify algorithms that were likely to have false positives and genes that were likely to be the result of HGT. Algorithms that had an outsized number of taxa missing from an orthogroup (METHODS) were considered false positives and removed from downstream analysis of that orthogroup’s age. After trimming these outliers, genes that were in the 95^th^ percentile of inferred losses were flagged as being potential recent HGT events (i.e. horizontally transferred long after LECA). These potential HGT genes are an interesting set in themselves: 66% are from the Euk+Bacteria category, they are hugely enriched for metabolic genes (gProfiler p-value=9.08e^−116^), and several are associated with human diseases.

We found that, as expected, algorithms in the “old” group tended to commit more false positive errors, and algorithms in the “young” group committed more false negative errors (Figure 5). Because PhylomeDB was used as a basis for identifying false negatives, its false negative rate could not be quantified.

**Figure 5.**
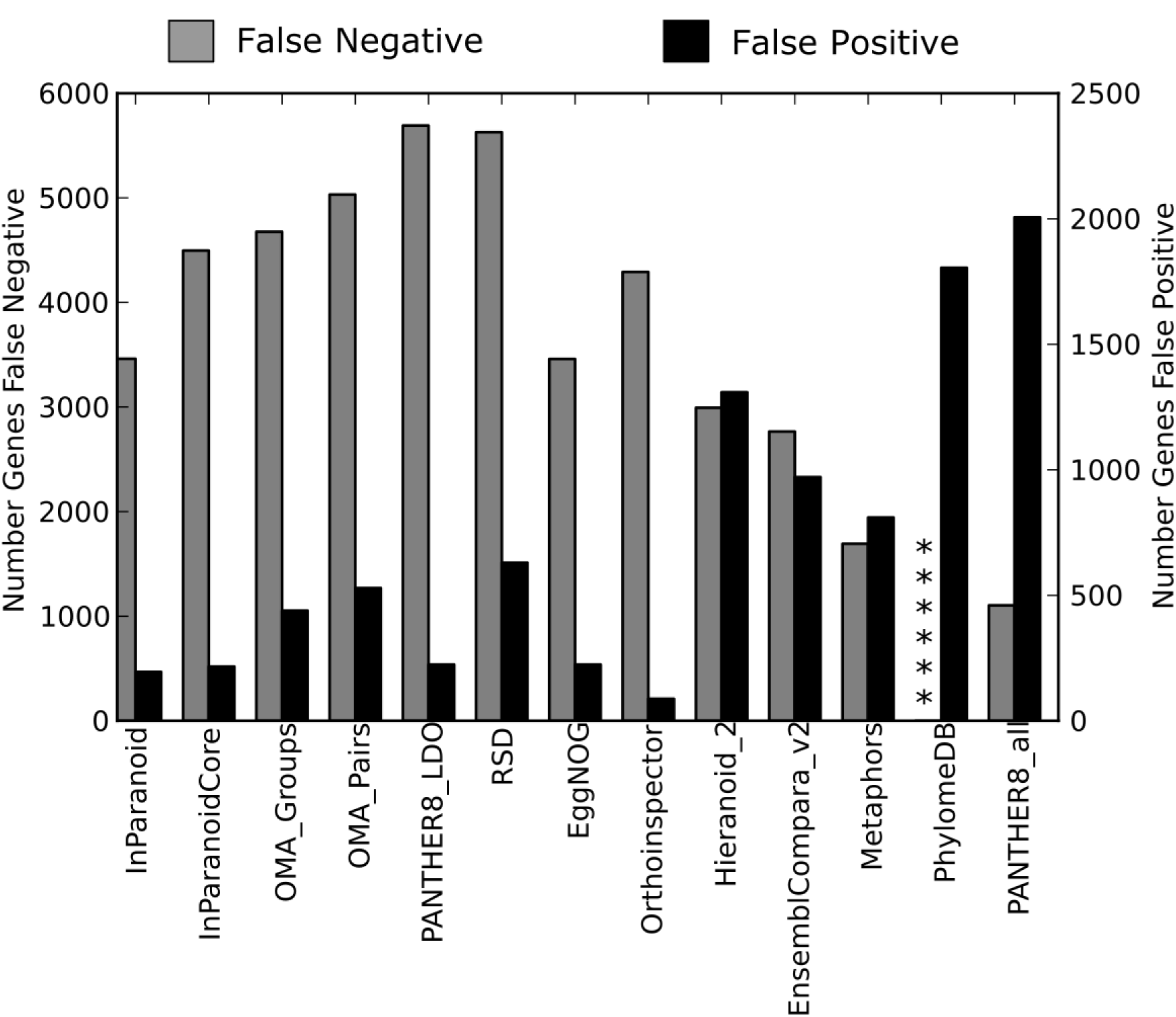
Errors committed by different algorithms. False positives and negative are defined in the text. PhylomeDB was used as a standard for false negative, so its false negative count could not be determined.

### Consensus

These analyses suggested a way to more robustly estimate consensus gene ages and to calculate a posterior distribution over the estimate. We used the methods described above to identify algorithms that may have committed false positive or false negative errors and then removed these algorithms from consideration on a per-protein basis. After doing so, we generated consensus tables based on the remaining algorithms for the human proteome and for a number of other model eukaryotes (Table 1), and we make these tables available (WEBSITE). Because our tree is best sampled within the opisthokonts (fungi, animals, and closely related protists), we restricted our analyses to this lineage. These tables contain a consensus age category for each protein based on the mode age call of non-trimmed algorithms. Older genes were found to be involved in key components of cell biology. Genes in the Euk+Bac group were found to be highly enriched for mitochondrial function, and genes that date back to the Euk_Archaea node were enriched for translational machinery, as has been shown previously [8,9]. Many of these older genes are also associated with hereditary diseases that represent a deficiency in a cell function associated with that evolutionary epoch. For instance, the cytoskeletal system and cilium date to LECA [9], and genes in this age category are enriched for diseases affecting the cilium, such as primary ciliary dyskinesia and Bardet-Biedl syndrome (Figure 6).

**Figure 6.**
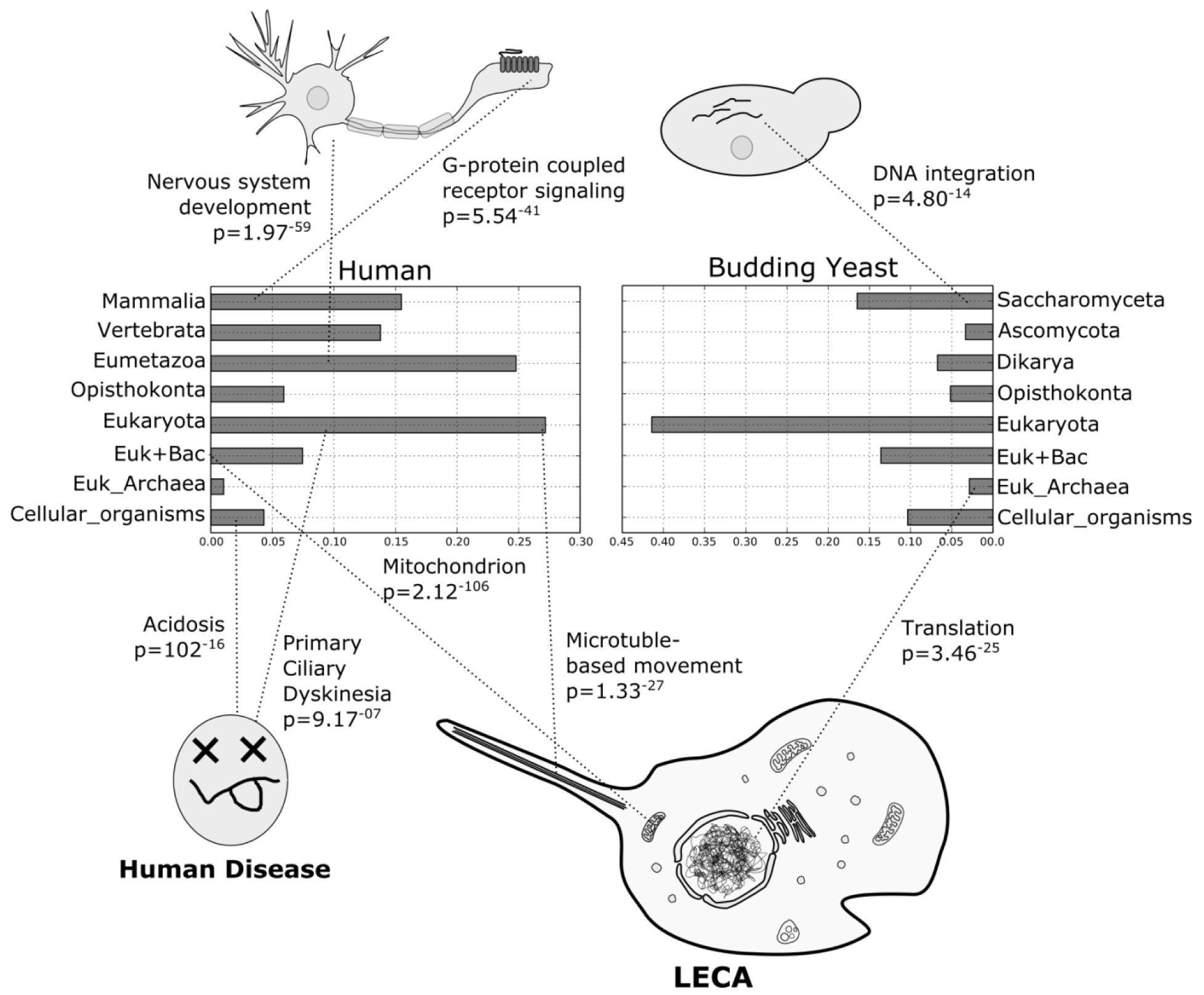
Enrichment of gene ontology terms and human disease terms (OMIM) in the different consensus age classes for human and budding yeast (*Saccharomyces cerevisiae*). The distribution of age classes are shown for each species. Older genes tend to be enriched for core cellular machinery and heritable diseases. Newer genes are associated with lineage-specific function, such as nervous system development and olfaction (*via* G-protein coupled receptors) in mammals, and DNA integration in yeast. P-values are dervied from g:Profiler [38].

These enrichment terms are derived from the point estimates of consensus ages, but we also provide other data that can be used to propagate uncertainty to downstream analyses. For each gene, the distribution over age-calls from the non-trimmed algorithms is given, as well as the number of contributing algorithms and the entropy of the age call distribution. 87% of human proteins had at least 5 algorithms contributing after trimming, and 59% had at least 10 out of a total of 13 original algorithms. In addition, the tables contain information on whether the protein was flagged as being a potential horizontal gene transfer event. Finally, we include the node error and bimodality statistics, both of which are measures of uncertainty that reference the reference species tree.

We note that in several cases we have made *ad hoc* decisions during the building of the consensus. For instance, algorithms were flagged as false positives if the number of taxa inferred to have lost the orthogroup was two standard deviations above the mean of all algorithms. These decisions were informed by the underlying distributions of values. Nevertheless, we supply the source data files, scripts used for these analyses, as well as interactive iPython notebooks, and we invite researches to explore and change parameters if they desire (https://github.com/marcottelab/Gene-Ages).

### Error Propagation

How can our error annotations be used in downstream analyses? Here we give an example of a simple stability analysis for gene ontology enrichment that uses these error terms. It has previously been shown that eukaryotic genes vertically acquired from Archaea are enriched for translation and RNA processing, whereas genes acquired horizontally from bacteria at the root of eukaryotes are enriched for metabolic processes (Figure 6, [8,9]). This conclusion relies on functional term enrichment, but what is the effect of different sources of error on these sorts of enrichment analyses? To investigate the robustness of this conclusion to different sources of error, we used the program g:Profiler [38] to perform functional enrichment analysis on the two age classes “Euk_Archaea” and “Euk+Bacteria” after filtering the datasets at varying levels of stringency (Figure 7A). We found that removing genes that were flagged as a possible late HGT event had a strong effect on the average p-values of functional annotation terms in the Euk+Bacteria age class but not the Euk_Archaea class (Figure 7B). This may be due to these genes being more commonly lost or to many bacterial genes being more recent HGT events (and hence being filtered out). The latter possibility would mean that many genes in this age category could be misidentified as being present in LECA, so these genes are good candidates for manual curation. Notably, filtering on different error terms can increase or decrease the significance of different terms, and, depending on the filtering strategy, the significance ranking of terms can be switched (Figure 7C and D). Analyses that rely on smaller test-sets of genes are likely to be much more strongly affected that these proteome-wide searches.

**Figure 7.**
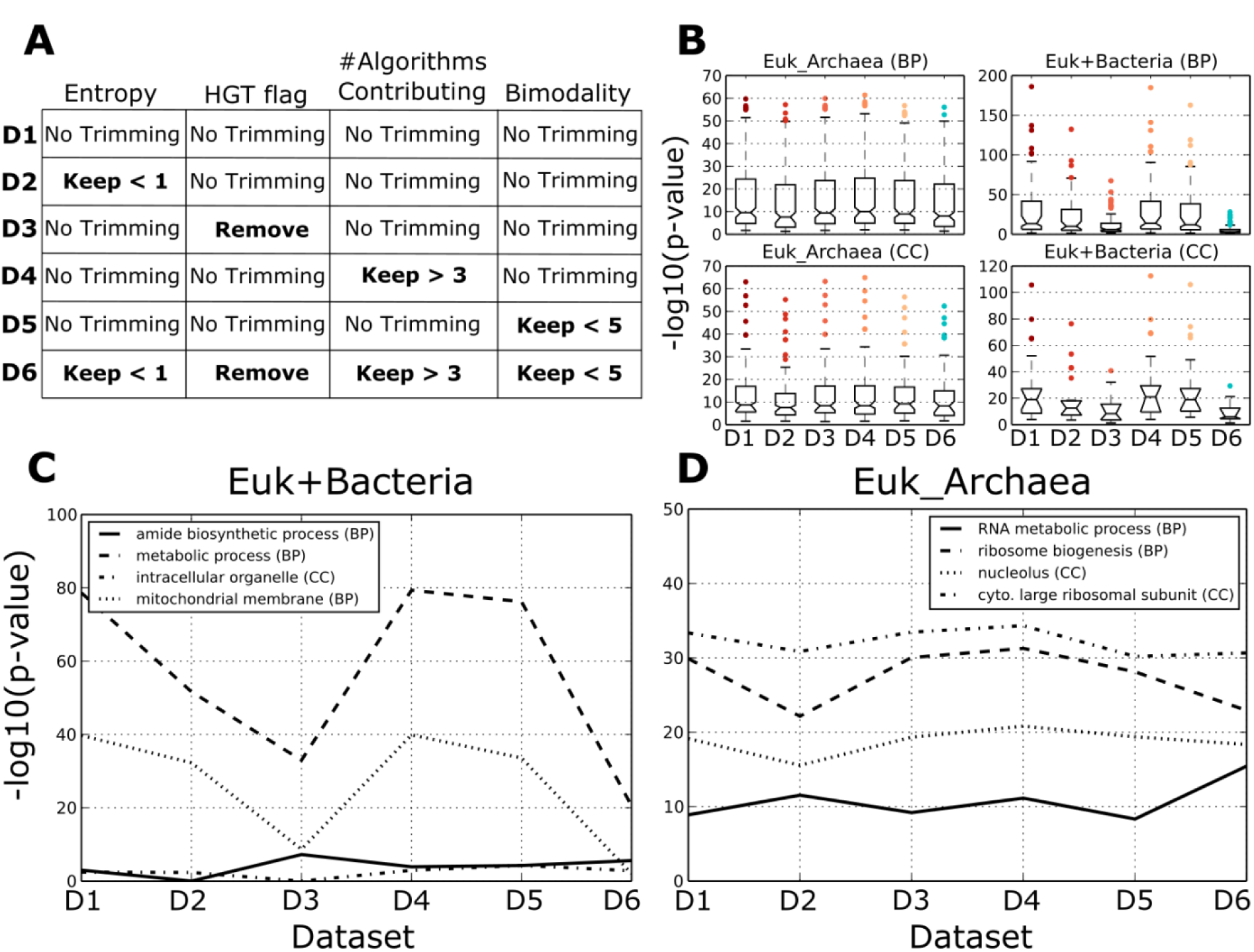
Effect of filtering on functional term enrichment analysis. (A) Datasets 1-6 were trimmed based on four sources of error: entropy of age-calls, whether the genes were flagged as potential horizontal gene transfer (HGT) events, the number of algorithms contributing to the final age call (after filtering algorithms, as described in Methods and Results), and the polarization of each gene. Dataset 6 was filtered on all four criteria (B) Negative log_10_ p-values for the five datasets for two age categories (Euk_Archaea, and Euk+Bacteria) and two gene ontology terms (Biological Process, and Cellular Compartment). (C-D) Negative log_10_ p-values across datasets for eight functional terms in the two age categories. These terms show the variety of ways that significance can be affected by filtering.

## Discussion

Most studies of gene age use a single point estimate arising from one of a variety of methods. Given our analysis of some the most popular orthology inference algorithms, we find that point estimates of gene age will be wrong for (at least) thousands of genes in a human-sized proteome (Figure 4). More troubling is the fact that algorithms appear to fall into two classes, each of which presumably has a systematic bias towards either false positives (“old group”) or false negatives (“young group”). This systematic bias happens on a per-gene basis, meaning that simple voting methods will not be able to resolve conflicts. Even with the ideal sampling of algorithms, which we approximate here by exploring a wide diversity of popular algorithms, the effective voting population will still drop to two on highly polarized genes.

Many areas of computational biology have faced a similar problem, namely, the need to keep track of error in several components of a workflow, and to correctly propagate this error through the whole analysis [39]. One illustrative example is multiple sequence alignment and phylogenetic inference. The former is a necessary precursor to the former, and each involves estimation error. Methods have been developed to infer the posterior distributions of both steps simultaneously [40], which is computationally intractable for all but the smallest datasets, or to perform each step iteratively in a maximum likelihood framework [41]. We argue that, eventually, such steps will have to be taken with orthology inference and gene-age estimation. Using a point estimate at each step in the analysis makes the assumption that each inference step has no uncertainty associated with it, which we can clearly reject in the case of gene-age estimation.

Some methods for probabilistic orthology inference do exist [21]. These use gene tree models with free parameters for gene duplication, loss, and sometimes HGT, which then contribute to the likelihood along with the multiple sequence alignment. However, these methods are in their infancy, and not usually scalable to large datasets or widely used. In the meantime, it is important to have an understanding of common sources of error in gene-age estimation. We provide that information along with consensus age calls for a variety of model organisms so that researchers can incorporate error propagation into their analyses in a way that is appropriate to their question of interest.

Several error terms are likely to be important for a broad range of analyses. The first and most straightforward is the entropy of the age-call estimate after filtering false positives and negatives. This statistic gives a quick idea of how certain an age-call is, with higher entropies being less certain. It is defined with reference to our age categories, so if researchers need to use other age categories, they must use the node age of the gene, which we also provide (Methods). HGT events are also likely to affect some datasets, especially when genes originating in Bacteria are involved (Figure 6). A large number of eukaryotic genes are likely transfers from Bacteria [8], but these may have been transferred at any point on the phylogeny. We define one age category, Euk+Bacteria, to describe all genes transferred before LECA, with later transfers hopefully being caught by our flag. If researchers are primarily interested in HGT, we suggest a much fuller analysis, as our simple method is likely to miss many HGT events. Finally, the bimodality of the age-call between “young” and “old” algorithm types is a key statistic. The systematic biases in the different algorithm types mean that many datasets will be radically different and difficult to compare, and it may account for some of the differences between studies of ancient gene repertoires that used either graph or tree-based methods. Genes that are highly polarized are good candidates for manual curation, because it is unlikely that any *ad hoc* algorithm will differ substantially enough from those we sampled here to be decisive.

Although we have characterized only two components of a typical computational biology workflow, orthology inference and gene-age estimation, it would be ideal to characterize error distributions for all the steps in an analysis, which has not been done with gene age data to our knowledge (but see [42] for an interesting example on gene-expression data, and [39] for a general review). The datasets we provide here will hopefully help guide future research efforts aimed at a more formal, probabilistic way to handle error in gene-age estimation, perhaps even in the context of an entire workflow. Until such methods are available, we advocate using our error annotations or a similar analysis in any study incorporating gene-age data.

## Methods

### Data Collection and Availability

15 algorithms have submitted their estimates on 66 reference proteomes (http://www.ebi.ac.uk/reference_proteomes) to QFO’s benchmarking tool (http://orthology.benchmarkservice.org/cgi-bin/gateway.pl). We omitted two of these because they either did not have full taxon coverage (RBH), or their results were so different from all the others that it dominated the variance in all downstream analyses (OMA_GETHOGS). Pairwise orthology calls for the 13 remaining algorithms were downloaded from the Quest for Orthologs benchmarking website [20]. These pairwise calls were converted into tables for each gene, which were then used for subsequent analyses. The reference species tree was downloaded from SwissTree [33] on 06/15/2015 (ftp://ftp.lausanne.isb-sib.ch/pub/databases/SwissTree/speciestree.nhx) and was pruned to match the taxa in the Quest for Orthologs reference proteomes (http://www.ebi.ac.uk/reference_proteomes). Custom programs were written to perform the analyses below, and these are publicly available, as are iPython notebooks used for plotting. These, and the datasets supporting the conclusions in this article are available on GitHub (https://github.com/marcottelab/Gene-Ages) with the following commit id (c1a2862fa894d7da4ccdf3fb8001e1b6b226bd09). Scripts relied heavily on the python packages dendropy [43], BioPython [44], and pandas [45].

### Protein Age Calls

All protein ages are referenced to the species tree obtained from SwissTree. The age of a protein is calculated on the species tree by finding the MRCA node of the taxa that have orthologs of that protein. This node is the “node age,” and the age group it falls into is the “binned age.” The binned ages conform to the interior labels given by SwissTree, with the exception of the Euk+Bac age category, which is not phylogenetic, but rather consists of proteins that are present in Bacteria and Eukaryota (and would thus normally be assigned to the oldest age class), but not in Archaea.

We calculate several measures of error amongst algorithms. First is an error statistic called “node error” based on the node age calls. Node error is the average number of branches (patristic distance) between the age calls any two algorithms. A similar measure was used to calculate the distance tree in Figure 1. The average patristic distance between age-calls for each pair of algorithms was used as input for a heuristic search in PAUP [46]. Next, because algorithms fall roughly into two groups (“old” and “young”), we calculate the “bimodality” of each protein. This is the difference between the average within group (“old” and “young”) node error and the average between group node error. Note that, although we call this statistic simply “bimodality,” it captures not just the bimodal nature of the age calls, but how different the two peak ages are. Thus the proteins with the highest bimodality score are those for which all the “young” algorithms call one age, all the “old” algorithms call a different age, and these ages are very far apart on the tree (Figure 3).

### Filtering False Positives and Negatives

Before calculating a consensus, we flag algorithms that may have committed false positive or false negative errors on a per-gene basis. These algorithms are then removed from consideration of that gene’s age. False positives are orthology calls that are substantially more distant than orthology calls by other algorithms, and have the effect of driving age deeper in the tree. These are found as follows. For each algorithm and each protein: 1.) the node age is calculated 2.) the number of taxa in the species tree descended from this node is found 3.) The number of taxa containing orthologs of the focal protein is subtracted from the number of descendant taxa. This number is the number of taxa without the orthogroup that are descended from an ancestor that putatively had the orthogroup, and is therefore proportional, but not identical, to the number of inferred losses of the orthogroup. For each algorithm and each protein, if this number is two standard deviations above the pooled algorithm mean for the focal protein, that algorithm’s age call is considered a false positive and is thrown out.

False negatives are cases where an algorithm fails to make an orthology call, driving the inferred age to shallower nodes in the species tree. We identify one possible cause of this, which we call “over-splitting.” This is when a group of co-orthologs is not correctly recognized by an algorithm and only one or a few of its members are found as orthologs to a more distant species, while the others are split off into their own orthogroups. The members that are split off would then be called at an incorrectly young age. To identify these errors, we used PhylomeDB’s [32] orthogroups as a standard. For each protein and each algorithm (except for PhylomeDB), if the focal algorithm called a younger age than PhylomeDB and a co-ortholog of the focal protein could be found where the focal algorithm called the same node age as PhylomeDB did on the focal protein, then this algorithm was considered to be over-splitting the focal protein, and was not considered in this protein’s age call. This error calculation was not performed on proteins where PhylomeDB was flagged as a false positive.

### Consensus Ages

We generated consensus binned ages after removing algorithms flagged with false positives and negatives as described above. The number of algorithms favoring each binned age is counted and then normalized by the number of contributing algorithms to give a distribution over age calls. For subsequent analyses, we used the mode of this distribution as the consensus age.

## Acknowledgements

We would especially like to acknowledge the Quest for Orthologs consortium and those who contributed their algorithms to the benchmarking tool for making their data freely available. B.J.L was funded by NIH fellowship 1F32GM112504-01A1. E.M.M. acknowledges funding from the NIH, NSF, CPRIT, ARO (61789-MA-MUR), and Welch Foundation (F1515).

